# Decoding Imagined Musical Pitch from Human Scalp Electroencephalograms

**DOI:** 10.1101/2022.06.13.495849

**Authors:** Miyoung Chung, Taehyung Kim, Eunju Jeong, Chun-Kee Chung, June-Sic Kim, Oh-Sang Kwon, Sung-Phil Kim

## Abstract

Brain-computer interfaces (BCIs) can restore the functions of communication and control in people with paralysis. In addition to the currently proven functions restored by BCIs, it would enrich life if one could regain a function of musical activity. However, it remains largely unknown whether it is feasible to decode imagined musical information directly from neural activity. Among various musical information, this study aimed to decode pitch information directly from scalp electroencephalography (EEG). Twenty healthy participants performed a task to imagine one of the seven musical pitches (C4 – B4) randomly. To find EEG features for pitch imagination, we took two approaches: exploring multi-band spectral power at individual channels (IC); and exploring power differences between bilaterally symmetric channels (DC). We classified these features into the seven pitch classes using various types of classifiers. The selected spectral power features revealed marked contrasts between left and right hemispheres, between low-, (<13 Hz) and high-frequency (> 13 Hz) bands, and between frontal and parietal areas. The best classification performance for seven pitches was obtained using the IC feature and SVM with the average accuracy of 35.68±7.47% (max. 50%) and the average information transfer rate (ITR) of 0.37±0.22 bits/sec. Yet, when we decoded a different number of classes (*K* = 2 ∼ 6) by grouping adjacent pitches, ITR was similar across *K* as well as between IC and DC features, suggesting efficiency of DC features. This study would be the first to demonstrate the feasibility of decoding imagined musical pitch directly from human EEG.

## I. Introduction

Connecting our mind to the world is fascinating when just hearing it, which has been struggled for long to develop fine systems. Thanks to recent advances in neurotechnology, active brain-computer interfaces (BCIs) have proven to restore brain functions by materializing brain activity patterns entrained from an imagery of certain tasks [1]. Motor imagery based BCIs (MI-BCIs) are one of the best-known active BCIs with its functional effect of moving a device using brain activity patterns elicited by imagining movements [1-2]. Especially, MI-BCI using electroencephalogram (EEG) is proven to help patients suffering from post-stroke syndrome by targeting damaged neuronal circuits and recovering motor function by training based on neuronal plasticity [2-3]. Those patients also suffer from post-stroke cognitive impairment (PSCI), including language, attention, and memory [4]. PSCIs were coped with traditional therapy, however, nowadays BCI-based training methods are introduced and prospected to have a promising effect on reducing cognitive anomalies [2,5].

A deficit of musical ability can also result from PSCIs, expressed as acquired amusia (AA) in patients who experienced a brain lesion with predominance in the right hemisphere by stroke, and restoring it seems crucial for cognitive functions extensively [6-7]. In addition to the enjoyable aspect of music, musical ability is known to be comprehensively correlated with non-musical cognitive functions, such as language, intelligence, memory, and attention [7]. Furthermore, musical ability is more intrinsic than language in terms of brain functions since patients with a severe level of dementia who lost language ability still possess musical ability, which postulates the possibility of a new communication channel for those who cannot operate adequate linguistic act [8]. Therefore, the rehabilitation of musical ability would derive positive effects on non-musical cognitive functions and neurological disorders [7].

One of the key musical factors affecting all-inclusive musical ability is pitch, defined as the feature of auditory sensation ordering from “low” to “high” [9]. The importance of pitch in musical ability has been revealed by studies of amusia (i.e., tone-deafness). There are broadly two types of amusia; acquired amusia (AA) and congenital amusia (CA). AA is manifested from brain damages caused by neurological disorders such as stroke, especially found in the right hemisphere over ventral and dorsal connectivity between frontal and parietal areas, which reportedly underlies pitch processing [6,10]. CA is an inherent disorder of comprehensive musical ability, and a study of patients with CA revealed that a scarcity of the ability to detect pitch changes was dominant among various musical ability measures, showing the priority of pitch processing [11]. Moreover, patients with CA exhibit the deformity at right frontotemporal cortical networks in the early development stage [12-13]. Recent studies have demonstrated that right dorsal connectivity was the key to AA recovery, and that a longitudinal pitch and melody discrimination training showed a potential to enhance the musical ability of people with amusia even if it is inherited [6, 14-15]. Accordingly, it can be conjectured that musical ability may be augmented by training pitch processing capacity.

The rehabilitation of musical communication functions via active BCIs would rely on the possibility of decoding pitch imagery-inducing brain activity patterns and training pitch imagery for BCI use [1-2]. Moreover, deriving motor-related strategies such as imagining the production of pitch by an instrument or singing would be effective for BCI training [7]. Such strategies have been employed by recent speech BCI studies. Willett *et al*. realized a high-performance mental typewriter by decoding handwriting imagery inducing motor cortical activity via intracortical BCIs [16]. Also, Moses and colleagues allowed patients with anarthria to type a sentence in real-time by decoding words from sensorimotor cortical activity acquired from subdural multi-channel electrodes [17]. Music and language are known to share neural pathways – both amusia and aphasia are related to dysfunctions of the right ventral stream [6]. Also, music and language share neural substrates including bilateral precentral gyrus (PCG) and superior temporal plane related to semantic and melody processing [18]. Especially, pitch processing is the most common between music and language, as pitch is used to construct intonation in non-tonal language, and semantic differences in a tonal language, and melody in music [19-20]. Thus, by tracking the footprint of speech BCIs, it can be hypothesized that pitch imagery based BCIs would be the cornerstone of building the track.

Many efforts have been devoted to enable musical communications via BCIs. A set of musical user interfaces based on BCIs were developed to allow the users to choose a musical property by indirect brain activity induced by sensory stimuli or mental state changes. For instance, a reactive BCI allowed the users to choose musical notes by selectively paying attention to corresponding visual stimuli. Also, the users controlled musical events such as sound intensity by modulating the intensity of attention via a BCI. Other BCIs enabled the choice of musical modules generating musical passage modeled from EEG frequency band power [21-23]. These studies do not harness musically or auditory relevant neural activity but use rather irrelevant brain activity to enable the selection of musical events.

Meanwhile, some closed-loop BCI studies have attempted to enhance other functions via musical stimuli: e.g., treating depression in the elderly by choosing the speed and loudness of the music according to arousal and valence levels of EEG signals [24]. Other studies decoded brain activity in response to musical stimuli into musical features but in a wider scope than just focusing on pitch. For example, a study of tonal hierarchy decoded two classes of pitch in a tonal relationship such as in-key/out-of-key, tonic/dominant, or minor 2nd/augmented information using multivariate pattern analysis of magnetoencephalograms (MEG) [25]. Also, Schaefer *et al*. decoded 7 segments of classical and contemporary music from the temporal pattern of EEG, and Sakamoto *et al*. decoded the contextual pitch information by classifying the position where the same pitch (440Hz) was when presented with a lower pitch (110Hz) or higher pitch (1760Hz) using connectivity features of EEG [26-27]. Despite all these efforts, no research has yet been conducted to decode individual musical pitches directly from brain signals. Therefore, the feasibility of decoding pitch on a musical scale from brain activity is still in veil.

Understanding the neural representation of pitch is crucial to decoding individual pitch information, guiding toward an effective strategy to find relevant brain activity features. Despite the lack of known neural mechanisms of pitch imagery, capitalizing on the perceptual processing of pitch will be helpful as the imagery and perception of sound is known to share common neural networks including the secondary motor area and dorsal premotor cortex [28]. Moreover, studies on temporal patterns of high-gamma oscillations in EEG and electrocorticograms (ECoG) showed that the perception and imagery of music shared activated regions but in different temporal orders; for example, frontal region was activated first for imagery while temporal or sensory cortex first for perception though these regions were all shared between perception and imagery [29-30]. Although neural processing of single musical pitch information in humans still needs further investigations, right lateralization is reported to be an essential attribute in pitch processing. Especially, patients with right lateral Heschl’s Gyrus (HG) resection exhibited a shortfall in the detection of pitch change direction, which supports a hypothesis that right HG plays the role of a ‘pitch centre’ [10, 31]. Besides, the right temporal and frontal cortices were involved in melody processing, with more involvement of the temporal cortex when an active pitch memory task was carried out [32]. An account of right hemisphere lateralization to musical pitch suggests a temporal sensitivity difference of information processing between the left and right hemispheres, with a longer temporal window in the right hemisphere resulting in a higher spectral resolution. As such, musical sounds containing more spectral features than linguistic sounds may be better processed by the right hemisphere [33]. In contrast, other studies suggested that the local pitch contour was related to the left posterior hemisphere [34-35]. Those studies imply the necessity of broader inspection along with right lateralized neural responses to musical pitch compared to the left. Furthermore, spectral characteristics of neural responses would be related to pitch perception as the auditory neural pathway processes spectro-temporal structures of sound information [10, 31].

This study aims to investigate the feasibility of decoding musical pitch imagery information directly from the human brain activity, especially from electroencephalogram (EEG). To find pitch-related spectro-temporal features from EEG, we embrace a comprehensive approach to explore all possible EEG features from the bilateral hemispheres, as well as to capitalize on the property of hemispheric asymmetry for musical pitch processing by extracting differences between paired channels of EEG across hemispheres. Specifically, we aim to decode seven musical pitches (C4, D4, E4, F4, G4, A4, and B4) from EEG features when people attempt humming of each pitch one a single trial basis. We focus on the spectral feature space with five frequency bands: delta, theta, alpha, beta, and gamma bands. We devise a heuristic and automatic method to select features, considering EEG spectral power differences among all pitch pairs. Here, we compare various classifiers, including Naïve Bayes’ classifier, multi-class Support Vector Machine (SVM), Linear Discrimination Analysis (LDA), XGBoost, and Long Short-Term Memory (LSTM) models, to choose the most appropriate model for our purpose. We also compare decoding performance between two participant groups, one with musical training and the other without it, to examine the potential effect of musical training. We anticipate that unveiling the feasibility to decode the imagined musical pitch from brain activity will be a step forward on the road to building a cornerstone to realize a music BCI for the restoration of musical communications.

## II. Methods

### A. Subjects

21 participants were recruited for this study. Among them, those who received formal musical training less than 3 years were allocated to the non-trained (NT) group. Those who received more than 3-years of musical training and passed a pitch detection ability test conducted in our experiment (see section II.B for more details) were allocated to the musically-trained (MT) group. As a result, 10 subjects were allocated to the MT group (5 females, average age of 24.2 years old) and 10 to the NT (4 females, average age of 25.5 years old), where 1 subject with musical training experience failed to pass the test and consequently was excluded from the study.

All subjects in the MT group were able to play the piano. No participant reported abnormalities in hearing and brain function. Written consent was obtained from all subjects before the experiment, and subjects were paid for their participation. This study was approved by the Institutional Review Board (IRB) of the Ulsan National Institute of Science and Technology (UNISTIRB-20-22-A).

### B. Stimuli

Auditory stimuli were composed of 7 pitches in the 4th-octave musical scale: C4, D4, E4, F4, G4, A4, and B4, including the international standard note of A4, to which most musicians tuned their instruments [36]. The frequency corresponding to each pitch was 261.63, 293.66, 329.63, 349.23, 392.00, 440.00, and 493.88 Hz, respectively. Each stimulus was created as a piano sound with a duration of 500 ms. The sound intensity was adjusted according to the individual comfort level of each subject, assessed by their verbal responses. All the stimuli were generated with the Logic Pro (Apple, Inc., USA).

A visual stimulus was designed as a piano keyboard image with the note names in Korean tagged on each of seven pitches. During the experiment, the visual stimulus was used to indicate the target pitch of the imagery task (see section C for more details).

### C. Experimental Task

Subjects performed two different tasks during the experiment: a perception task followed by an imagery task. In the perception task, subjects perceived 50 pitches presented in series with a 500-ms inter-stimulus interval (ISI) between successive presentations. Before pitch presentation, subjects were informed of a target pitch randomly selected from 7 pitches. Then, each of the 7 pitches was randomly presented for 50 times and subjects covertly counted the number of target pitch presentation. Afterward, subjects reported the count by pressing a keyboard. This round of perceiving 50 pitches and counting the target pitch was termed as a block. Subjects performed a total of 14 blocks, in which each pitch was selected as a target twice. We pseudo-randomized stimulus presentations such that each pitch was equally presented 100 times in total over 14 blocks. During the perception task, an image of the piano keyboard without any mark was displayed on the computer screen to help subjects concentrate on a musical scale. The performance of the perception task was used to determine a final MT group (see Section II.A). For those subjects with >3 years of musical training, if the number of correctly counted blocks exceeded 10, the subject was allocated to the MT group; otherwise, the subject was excluded from the experiment. This exclusion criterion did not apply to the subjects without musical training.

In the imagery task, subjects imagined the vocal production of a pitch on a musical scale (Fig. 1A). Again, there were 14 blocks in the task, where a block consisted of 50 trials of imagining a series of pitches randomly. At the beginning of each block, subjects initially listened to 7 musical pitches from C4 to B4 in sequence, which was designed to help subjects create a mental representation of the musical scale. Then, subjects conducted a random imagery task with the guidance of a visual cue displayed on the image of a piano keyboard. The visual cue was designed to turn one of the 7 keys to red for 500 ms followed by an ISI of 500 ms. In each trial, the visual cue appeared on one of the 7 pitches randomly and subjects were instructed to imagine the vocal production of the cued pitch by humming. Subjects performed 50 trials per block without any auditory stimulus. After the 50 trials of the imagery task, subjects were instructed to overtly hum the randomly selected 3 pitches, which were used to monitor how well they were performing the imagery task. Through 14 blocks, we acquired 100 trials for each pitch via pseudo-randomization. Subjects took a break after each block as much as they wanted but no longer than 2 minutes.

**Fig. 1.**
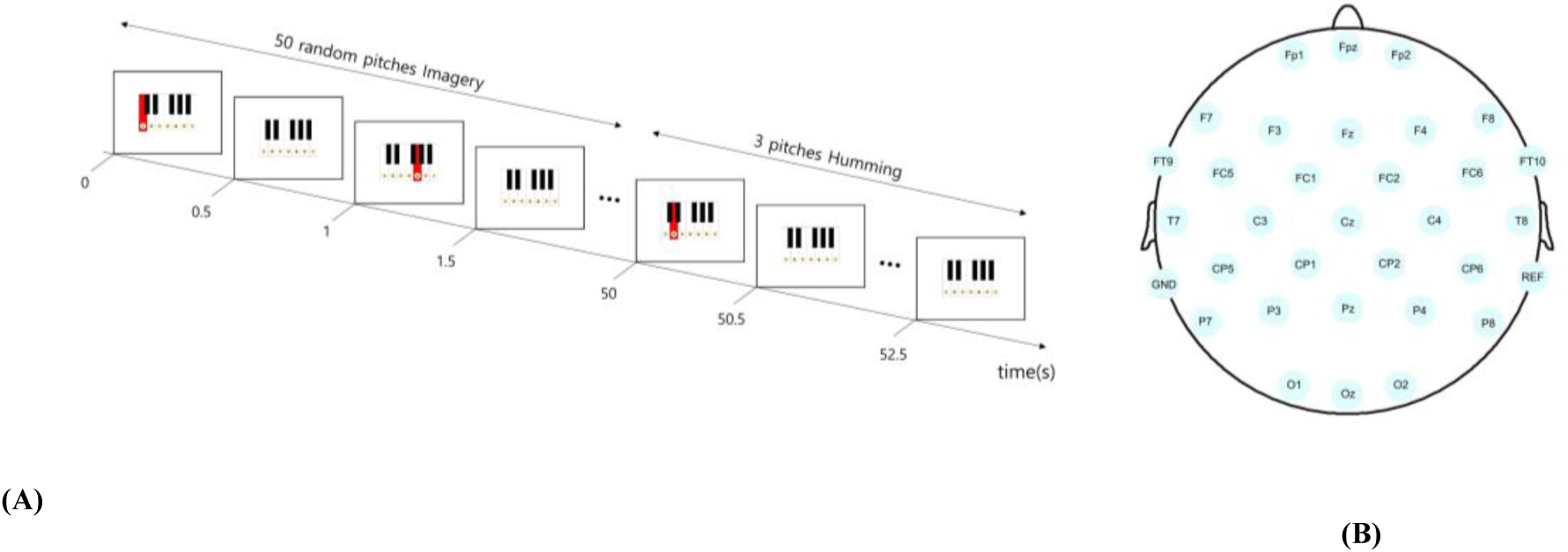
**(A) A pitch imagery task**. The pitch imagery task included 50 trials of pitch imagery. In each trial, a visual cue (red keyboard) appeared randomly on one of the seven notes (C4 – B4) for 0.5 s, cueing participants to imagine the corresponding pitch covertly. A 0.5-s inter-stimulus interval followed with no cued keyboard. After 50 trials, participants hummed overtly following visual cues randomly presented for 3 times. **(B) EEG channel Montage**. Thirty-one EEG electrodes were placed at the locations following the 10/20 international standard.

The experimental paradigm was implemented by MATLAB Psychtoolbox (Mathworks, Inc. MA, USA). The visual stimuli were presented on the LCD monitor of 1920×1080 resolution and the auditory stimuli were presented through earphones plugged into both ears. In this study, only the data from the imagery task were analyzed as the study’s goal was to decode imagined pitch from brain activity.

### D. EEG Data Acquisition and Processing

EEG signals were acquired by a commercially available EEG amplifier (acti-Champ, Brain Product GmbH, Germany) with 31 active wet electrodes (FP1, FPz, FP2, F7, F3, Fz, F4, F8, FC9, FC5, FC1, FC2, FC6, FC10, T7, C3, Cz, C4, T8, CP5, CP1, CP2, CP6, P7, P3, Pz, P4, P8, O1, Oz, and O2) at a 500-Hz sampling rate (Fig. 1B). Reference and ground electrodes were placed on the mastoids of the left and right ears, respectively. The impedance of most electrodes was kept below 10k-Ohm except for a few electrodes (3 in maximum), where any impedance did not exceed 20k-Ohm.

The preprocessing procedure of EEG signals was conducted as follows: 1) The EEG signal at each electrode was high-pass filtered above 1 Hz; 2) The line noise was removed using a notch filter at 60 Hz with a 2-Hz bandwidth; 3) The signal was band-pass filtered with a passband from 1 Hz to 50 Hz; 4) Bad channels were detected and removed using a correlation criterion as follows: at each channel, when low-pass filtered signals (<1-Hz) exhibited a cross-correlation lower than 0.4 with more than 70% of all other channels’ 1-Hz low-pass filtered signals, the channel was deemed to be a bad channel [37]; 5) The common average reference (CAR) technique removed a potential common noise component from the reference; and 6) Artifacts were further eliminated by the artifact subspace reconstruction (ASR) method with a cutoff of 30, reported to be appropriate to preserve brain activity and reject artifacts simultaneously [38].

The preprocessed EEG signals were then epoched from −200 to 1000 ms after stimulus onset. In each epoch, event-related spectral perturbation (ERSP) data was extracted as follows: First, the epoched signal at each channel was transformed to time-varying spectral power data via short-time Fourier transform (STFT) with a 100-ms sliding window and 90% overlap. By setting the number of points to the discrete Fourier transform to be computed as 512, the frequency resolution was 0.98 Hz. As the preprocessed EEG signals were filtered between 1 Hz and 50Hz, the number of spectral power values within this range (*f*) was 50. In addition, sliding windowing above yielded 111 time samples (*t*) within the epoch. Thus, the result of STFT for one trial was *X*_*STFT*_ ∈ *R*^*f*×*t*×*CH*^ = *R*^50×111×31^.

The baseline correction was performed for every frequency bin (here, the bin size = 1 Hz) by dividing all the spectral power values within that bin by the baseline mean, which was set as −200 ms to 0 ms. Then, the baseline-corrected values were transformed to the log scale. The baseline-corrected spectral power values were averaged across frequencies within each of the five frequency bands (delta: 1∼4 Hz, theta: 4∼8 Hz, alpha: 8∼13 Hz, beta: 13∼30 Hz, and gamma: 30∼50 Hz). The averaged spectral power values in each band were normalized over time within the epoch by the z-score method. As we intended to extract imagined pitch information guided by the cue (i.e., 0 to 800 ms after stimulus onset), we collected band power values after stimulus onset with 81 time points. As a result, a matrix *X*_*input*_ ∈ *R*^5×81×31^ from each trial was submitted to the subsequent feature extraction procedure. We assigned the class label to each trial according to the pitch information (a total of 7 classes). Considering real-time implementation of decoding in BCIs, we arranged the trials for each pitch class in a chronological order. Then, we included the first 80% of trials in a train set and the last 20% in a test set for each class. As such, there were 560 trials in the train set, *X*_*train*_ ∈ *R*^5×81×31×560^, and 140 trials in the test set, *X*_*test*_ ∈ *R*^5×81×31×140^, respectively.

### E. Feature Extraction

Using the train set, we explored spectral power features over time that were most distinguishable across the seven pitches for every frequency band and channel. This exploration was performed based on two different schemes: using spectral power values at individual channels (IC) and using spectral power value differences between bilateral channels (DC).

In the IC scheme, we examined the time courses of spectral power for different pitches in each frequency band at a given channel (e.g., see Fig. 2A-B). From the visual inspection of the time courses averaged over trials, we observed that different time courses crossed at a certain time point after stimulus onset, and then diverged. Such divergence of the time courses appeared to be bmaximal immediately after convergence. This pattern of time courses was observed in most bands and channels (Fig. S1). Based on these observations, we devised a metric to assess the divergence of the time courses according to pitch at each time instant as follows:

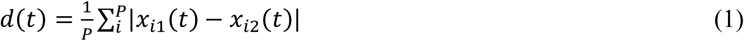

where *d*(*t*) is the mean of pairwise absolute differences between all pairs of 7 pitches in band power at time *t*, and *x*_*i*1_(t) and *x*_*i*2_(t) are averaged band power at time *t* of the *i*-th pair of pitches for *i* = 1, …, *P*, where *P* is the number of pitch pairs (*P* = 21). We first identified negative and positive peaks of *d*(*t*). Among these peaks, we inspected a pair of a negative peak and a positive peak, where the negative peak preceded the positive one, and selected the pair that showed the biggest difference between peaks, as it could reflect the crossing followed by divergence of the time courses that we observed. The selected peaks were used to determine the time segment for feature extraction, which corresponded to the full width at half maximum (FWHM) of the gap between negative and positive peak (see blue lines in Fig. 2.D-E). Then, a feature was extracted as an area under the time courses of spectral power within this time segment for every trial.

**Fig. 2.**
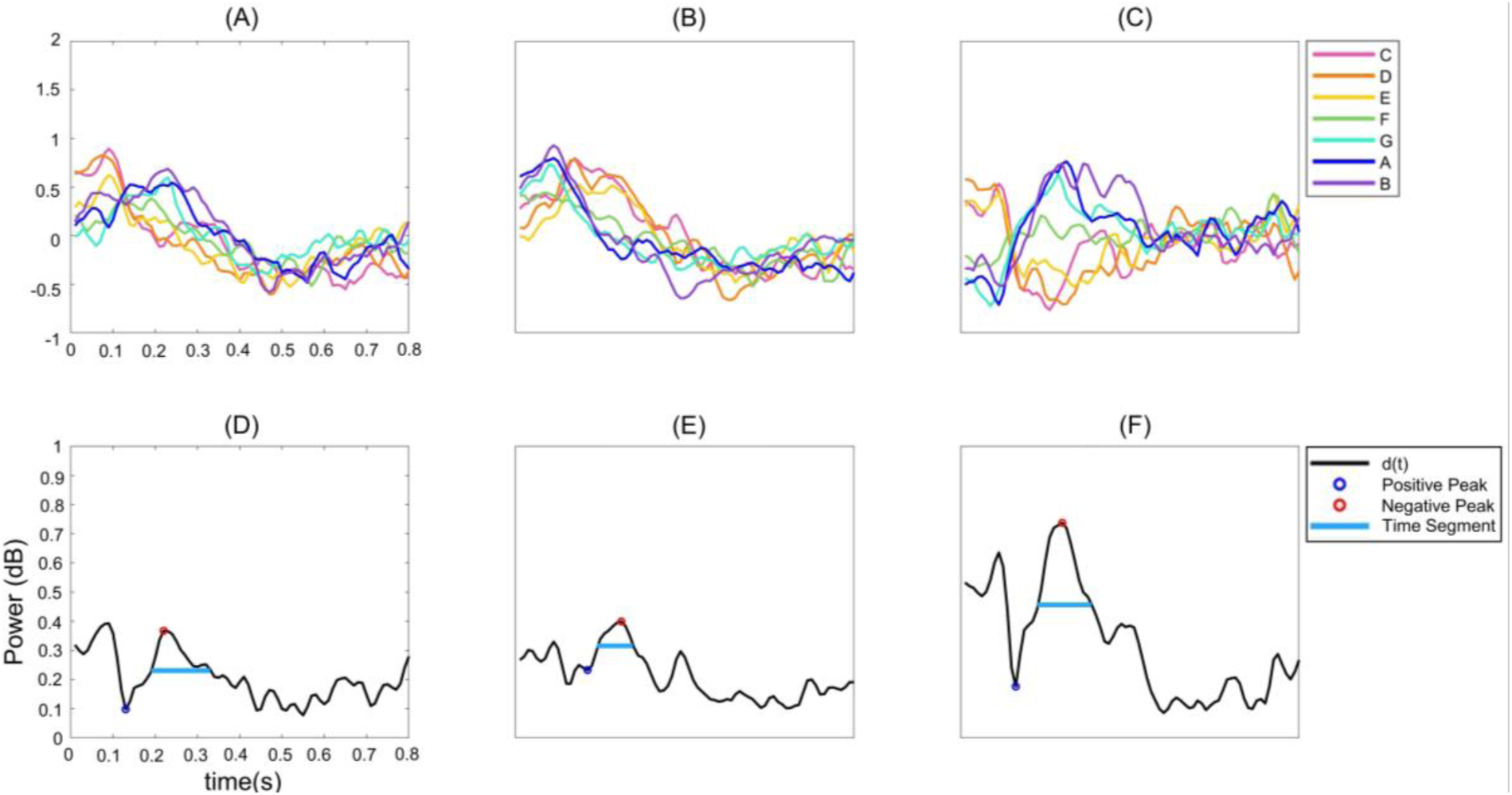
Feature extraction procedure. The time courses of the mean theta band power from the visual cue onset (0 s) to the end of an epoch (0.8 s) for each of the seven pitches (C4 – B4) are depicted for EEG channels of F3 (A) and F4 (B), respectively. Differences of the time courses between the two channels (F3-F4) are also depicted (C). The divergence of the time courses across pitches (*d*(*t*)) for each set of the time courses at F3 (D), F4 (E) and their difference (F) is depicted. The positive (blue circle) and negative (red circle) peaks used for time segment selection, and the resulted time segment (cyan line) are marked. Note that the power values at 0 s (A)∼(C) can vary as each time course is normalized independently.

In the DC scheme, we extracted features from hemispheric differences of band power based on our observation that the time courses of band power according to pitch exhibited opposite patterns between the left and right hemispheric channels (see Fig. 2. A-B). To capitalize on such contrast, we subtracted the band power of the right hemispheric channel from the left counterpart for each channel pair, where 13 bilaterally symmetric channel pairs were arranged as follows: Fp1-Fp2, F7-F8, F3-F4, FC9-FC10, FC5-FC6, FC1-FC2, T7-T8, C3-C4, CP5-CP6, CP1-CP2, P7-P8, P3-P4, and O1-O2. Consequently, the crossing and divergence of time courses of band power shown in individual channels became more pronounced, as demonstrated in Fig. 2C. From the time courses of band power, a set of features was extracted in the same way as the IC scheme using *d*(*t*) (Fig. 2F). The selected positive peaks of *d*(*t*) in the DC scheme showed larger magnitudes than those in the IC scheme (Fig. S2).

From 5 bands with 31 channels or 13 channel pairs, the size of the feature set was 155 through the IC scheme or 65 through the DC scheme. Then, we further selected features from each feature set (IC or DC) using the one-way ANOVA test that showed a significant difference among the 7 classes (*p* < 0.01). The number of selected features from each set was 123.9±16.6 with the IC scheme and 54.9±5.96 with the DC scheme on average across subjects.

### F. Decoding Model

This study compared several classification algorithms to decode the selected features into 7 pitch classes, including Naïve Bayes’ classifier, multi-class SVM, LDA, XGBoost, and LSTM. The performance of the classifiers was evaluated by test accuracy, the information transfer rate (ITR), and the diagonality of the confusion matrix.

The test accuracy was calculated by the ratio of the number of corrected trials to the total number of trials in the test set. ITR was calculated as follows [39]:

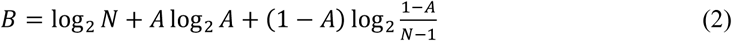

where *N* is the number of classes, and *A* is the test accuracy. We divided *B* by 0.8 sec, the length of an epoch, to obtain ITR in the unit of bit /sec.

The diagonality of a confusion matrix was also calculated to evaluate how close a misclassified pitch was to a true one. As the pitch classes were assigned in a linear fashion, misclassification into a pitch far from a true one could be regarded worse than misclassification into a closer pitch, which would represent the degree of confusion. For example, if the true class was C4, decoding as D4 would be considered less confusing than decoding as A4, although both were treated misclassification in terms of accuracy. Thus, we devised a simple method to assess the diagonality of a confusion matrix by correlating a confusion matrix with a semi-diagonal matrix, where the semi-diagonal matrix contained 2 in its diagonal entries and 1 in the entries adjacent to diagonal ones (Fig. S3). We calculated a 2-D correlation (*r*) between two matrices given by:

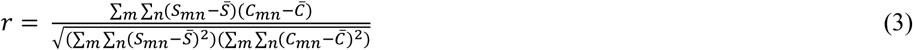

where *S* is the semi-diagonal matrix and *C* is a confusion matrix. *S*_*mn*_ (*C*_*mn*_) is an entry of *S* (*C*) at the *m*-th row and *n*-th column and 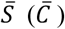 is the mean of all the entries in *S* (*C*).

Not only did we decode 7 individual pitches, but we also decoded a group of pitches to investigate whether decoding performance was increased if we reduced the number of classes. We grouped pitches into *K* classes (1<*K*<7) under the following condition: The groups must be chunked linearly according to the pitch height, pitches in each group must be adjacent to each other, and the number of pitches per group must be as balanced as possible. For example, with *K*=3, pitches could be grouped as CDE/FG/AB, CD/EFG/AB, or CD/EF/GAB, but not as CDG/EA/FB (pitches are not adjacent) or C/DEFG/AB (not the most balanced). For each value of *K*, we explored all possible ways of grouping pitches, and found the one with the highest classification accuracy with all classifiers.

## III. Results

In this section, we first present extracted features by the IC and DC schemes. Next, we present the decoding results for *K* classes (*K* = 1, 2, …, 7) using either the features extracted via the IC scheme (IC features) or the DC scheme (DC features). Thus, there will be 7 × 2 combinations of decoding outcomes. Note that we present the best decoding result among all possible ways of grouping 7 pitches into *K* classes when 1< *K* < 7. Lastly, we compare decoding outcomes between the MT and NT groups.

### A. Feature distribution

We explored the spatial distributions of the IC features and the DC features obtained from the training set as shown in Fig. 3 (showing examples of a representative subject (subject 11)). For the IC features, we observed systematic changes in the spatial distributions as the imagined pitch height changed (Fig. 3A). Frontal regions showing greater feature values gradually migrated from right to left hemispheres as the pitch height increased in the lower-frequency bands, including delta, theta and alpha bands. On the contrary, they migrated from left to right hemispheres as the pitch height increased in the higher-frequency bands, including beta and gamma bands. Similarly, temporoparietal regions showing greater feature values migrated from left to right hemispheres as the pitch height increased in the lower-frequency bands, which was seemingly reversed (from right to left) in the higher-frequency bands.

**Fig. 3.**
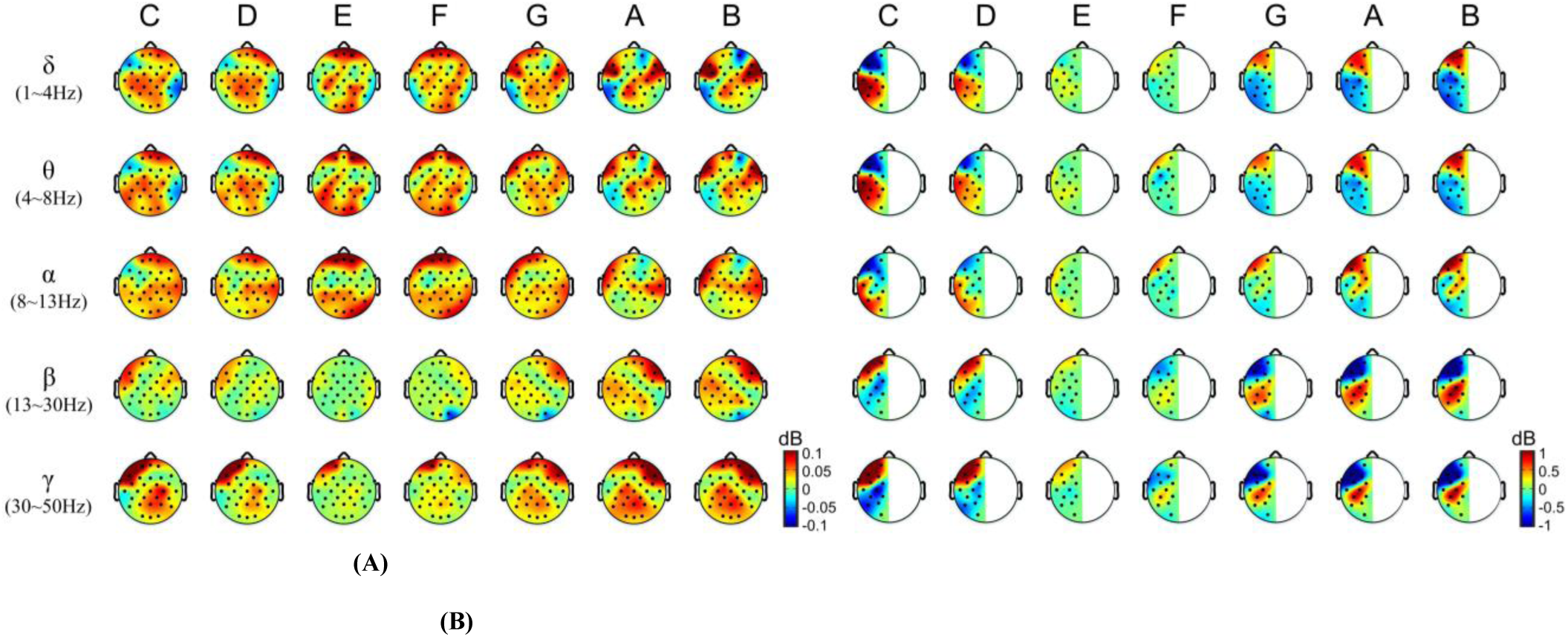
Topographic distributions of EEG spectral features. (A) Individual channel (IC) features, and (B) Difference between bilateral channel (DC) features. DC features are displayed on the left hemisphere for convenience.

The characteristics of the spatial distributions of features according to the pitch height was manifested more vividly with the DC features (Fig. 3B), which also dovetailed with the result in Fig. S2. Note that Fig. 3B depicts feature differences between hemispheres on the left hemisphere for visualiztion purpose, and a channel difference was calculated by subtracting right hemispheric feature values from left counterparts. We observed marked contrasts in variations of the DC features with the pitch height between frontal and temporoparietal, as well as between low-, and high-frequency bands, with a clear interaction between brain region and frequency. The DC feature values increased as the pitch height increased in the frontal region and the low-frequency bands (delta, theta and alpha) or in the temporoparietal region and the high-frequency bands (beta and gamma). In contrast, they decreased as the pitch height increased in the frontal region and the high-frequency bands or in the temporoparietal region and the low-frequency bands.

The spatial distributions of the selected IC or DC feartures largely remained consisitent across single trials of the training set (Fig. S4). Also, the spatial distributions of the features were more similar between adjacent pitches. Especially, the spatial distributions of the features appeared to be clustered into {C, D}, {E, F, G} and {A, B}. The selected feature distribution was averaged by pitch classes and depicted in Fig. S5 for all subjects.

### B. Decoding Individual Pitches

We first evaluated the decoding of individual pitches from the IC or DC features. We classified the features obtained from the test set into 7 classes using each of the five classifiers (see Table S1). Then, we compared classification accuracy and ITR among the classifiers and between the feature schemes (IC vs. DC) using the 2-way Scheirer-Ray-Hare (SRH) test. It showed no main effect of the classifier (p>0.05) but a significant main effect of the feature scheme (p<0.01). There was no interaction effect (p>0.05). A post-hoc analysis using the Kruskal-Wallis (KW) test revealed that the IC feature yielded better performance in accuracy and ITR than the DC feature (p < 0.05). The best decoding performance was achieved by using the IC features with SVM, resulting in the average accuracy of 35.68±7.47% (max. 50%) and the average ITR of 0.28±0.16 bits/sec (Fig. 4).

**Fig. 4.**
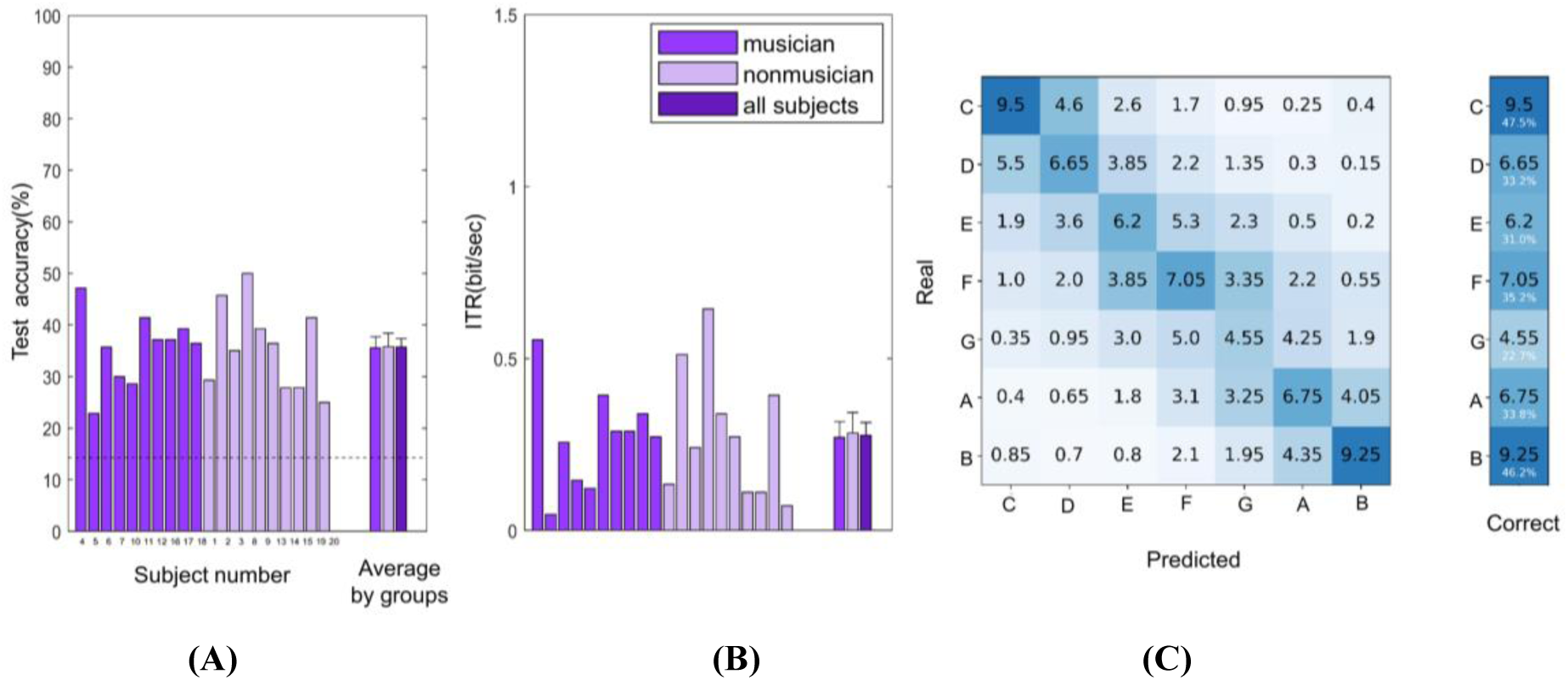
Decoding performance. The performance of decoding seven pitches from EEG obtained by the best combination of a feature set (IC features) and a classifier (SVM) is illustrated in terms of (A) Accuracy, (B) ITR, and (C) Confusion Matrix for individual subjects as well as subject-average. The MT and NT groups are displayed in different colors

We constructed the confusion matrix from the classification outcomes of the 5 classifiers for each the 2 feature schemes, respectively (Fig. S6-7). The SRH test showed the main effect of the classifiers (p<0.05), followed by the Dunn-Test showing that using LSTM yielded lower diagonality than using other classifiers (p < 0.05), (Fig. S8).

### C. Decoding Groups of Pitches

Classifications into *K* pitch classes were evaluated for each *K*, 1<*K*<7. Grouping of pitches showing the highest decoding accuracy averaged across the IC and DC features for each *K* was as follows: CD/E/F/G/A/B for *K*=6; CD/E/F/G/AB for *K*=5; CD/E/FG/AB for *K*=4; CD/EF/GAB for *K*=3; and CDE/FGAB for *K*=2.

For each *K*, we calculated accuracy and ITR using either the IC or DC features with each of the five classifiers (Table S1), and compared among the feature scheme and classifiers using the SRH test. For *K*=2 and *K*=5, the test showed a main effect of the classifier and the Dunn test revealed that LSTM yielded the highest accuracy (p<0.05). For *K*=3, the test showed a main effect of the feature scheme and the KW test revealed higher accuracy with the IC feature (p<0.05). For *K*=4, the test showed main effects of both classifier and feature scheme and post-hoc analyses revealed higher accuracy using LSTM and the IC feature (p<0.05). For *K*=6, no significant difference was found among all classifiers and feature schemes (p>0.05). There was no interaction effect for all *K* classes.

We selected the best combination of decoding models with the statistically optimized combination for each *K*. If there was no main effect of feature, we selected the DC feature as it required a smaller number of features than the IC feature to achieve the similar level of performance. Moreover, if there was no main effect of classifier, the best classifier was selected which could be most effectively implemented. For instance, the LSTM was not selected unless it showed significantly higher accuracy than others, as it takes a much longer computation time. The accuracy results from the best combinations were 84.07±12.34% (max. 96.43%) for 2 classes, 65.21±7.42% (max. 77.86%) for 3 classes, 58.18±10.75% (max. 80%) for 4 classes, 56.11±2.17% (max. 57.14%) for 5 classes, and 39.5±5.53% (max. 51.43%) for 6 classes, respectively (Fig. 5 top). All these accuracy values were significantly higher than corresponding chance levels, calculated as 1/*K* (t-test, *p* < 0.05).

**Fig. 5.**
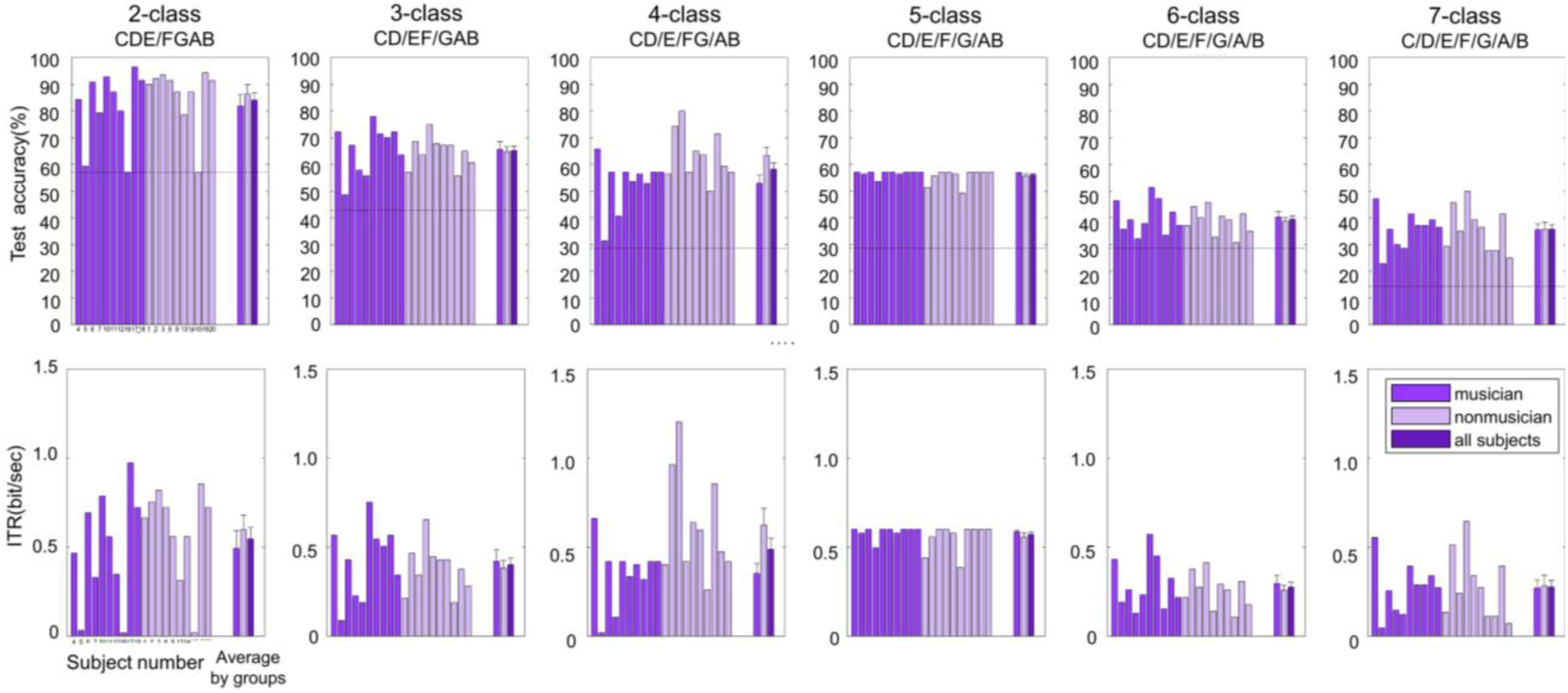
Decoding performance for a different number of classes of pitch. Decoding performance for *K* classes of pitch (*K* = 2, 3, …, 7) is illustrated in term of (top) Accuracy and (bottom) ITR. The optimal grouping of pitches for *K* classes is described over accuracy graphs. The best combination of feature and classifier is DC feature & LSTM for 2-, 4-, and 5-class, IC & LSTM for 3-class, and IC & SVM for 6-, and 7-class.

To compare the decoding outcomes among *K*, we evaluated ITR of selected decoding models for each *K* (Fig. 5 bottom). As ITR measures performance by taking the number of classes into account, it could reveal decoding performance more consistently across different number of classes. Firstly, for all the feature sets, there was no interaction between classification model and the number of classes (*K*) (SRH test, p>0.05) and the number of classes showed the main effect. The effect of *K* was tested for each feature types (Dunn test). As a result, the ITR for 5-class showed a relatively higher value. Specifically, with the IC feature, the ITR for 5-class was significantly higher than those for 2-, 6-, and 7-classes (p<0.05, Fig. S9A). With the DC feature, the ITR for 5-class was significantly higher than those for 3-, 4-, 6-, and 7-classes (p<0.05, Fig. S9B).

Furthermore, we visualized the feature distributions of the train set using t-SNE to investigate the feature distributions according to pitch groups for each *K* (Fig. S10). It showed that the DC features were distributed more linearly with pitch height. However, as the IC feature showed a more diverse distribution, chunked in various, the 7-class classification might be better with the IC feature.

### D. Comparison of Musically Trained and Non-trained Groups

We evaluated whether decoding performance was different between the MT and NT groups. We tested the results using the decoding models selected for each K as above (see Section III.C). The Dunn test revealed no significant difference in accuracy between the groups for all *K* classes (p>0.05). It also revealed no significant difference in ITR between the groups for all K classes (p> 0.05). Finally, no significant difference in the diagonality was found between the groups for all K classes (p>0.05).

## IV. Discussion

In this study, we decoded the pitch imagery information from EEG data with a feature extraction method of finding the most discriminable time segment of the temporal patterns of spectral power in every channel and frequency band. We designed two schemes for feature extraction, depending on whether features were extracted from individual channels (IC) or differences between channels (DC). In particular, differences were obtained between bilateral channels located symmetrically at each hemisphere, according to our observation that feature distributions were anti-symmetrical across hemispheres (Fig. 3). We used each of the IC or DC feature set to decode pitch using one of the 5 classifiers and selected the best combinations for each classification of *K* (2 ≤ *K* ≤ 7) classes in a statistically and computationally optimized way (Table S1). Classification accuracy was significantly higher than a chance level for every *K*, although it was not high enough to promise imminent applications to real-time BCIs. Between the feature sets, using the IC features was better in classification of multiple pitch groups (i.e., large *K*), whereas using the DC features was better in representing relatively higher or lower pitch (e.g., C4 or B4). However, when evaluating ITR, both types of features showed no difference for multiple pitch groups, suggesting that using the DC features was valid in representing pitch height information. Moreover, the results imply that the DC features would be more efficient than the IC features in terms of feature dimensionality, as using the DC features could lead to decoding performance close to that by using the IC features with only half of the number of features.

Exploring the feature distribution tracking the pitch height revealed noticeable countering distributions between 1) left vs. right hemispheres, 2) anterior vs. posterior areas, and 3) low vs. high frequency bands. Possible hypotheses for these contrasts can be posed with some neurological notions. The countering distribution between hemispheres (1) may be related to the temporal sensitivity difference between the hemispheres, making each hemisphere process pitch information with different levels of spectral resolutions [33]. Therefore, the relatively higher or lower frequency of pitch could probably form a bilateral alignment of features across the hemispheres. A potential neural substrate for the observed anterior-posterior contrast (2) can be conjectured by fronto-parietal networks related to pitch discrimination, though the source of distribution patterns according to pitch height is still in veil [35, 40]. The frequency-dependent distribution (3) may be related to the characteristics of different EEG oscillations tracking the acoustic properties of auditory stimuli, where delta to alpha oscillations reflected attentional and acoustic input variation in speech envelop for syllable or prosodic structures [41].

We further examined the commonality of the selected IC features amid subjects by counting the number of subjects from whom a feature was selected at each channel and frequency band (Fig. S11). Note that only the IC feature result was used here to scrutinize the whole brain distribution of the selected features. The features selected from most subjects were distributed over bilateral frontotemporal and parietal areas, especially in lower-frequency bands. These areas correspond to fronto-temporo-parietal networks related to pitch sensation recovery for AA and CA patients. Furthermore, the features in the low-frequency bands were frequently selected from lateral areas than medial areas, implying pitch information processing pathways over lateral areas. In the high-frequency bands, the selected features showed distributions in a more complicated manner. This may suggest that the DC features should be extracted by more meticulous inspection than simply subtracting powers between counterpart channels from the left to right hemispheres.

We examined five classification models including relatively more advanced models such LSTM and XGBoost with the expectation of superior performance. However, these models did not outperform other simpler models in this study. This may be related to insufficient amount of training samples for the advanced models [42]. Therefore, enlarging the training data size may be required to excel the performance of advanced models, which can increase the possibility of realizing a practical pitch-imagery BCI.

Despite our experimental design relying on the ability to imagine musical pitch, the MT and NT group difference was not found. EEG features representing pitch in a musical scale were evoked regardless of musical training. A possible reason might be the lack of musical contexts in the stimulus presentation [43]. Perhaps pitch imagery in a melodic context (e.g., in a song) might differentiate neural features of the MT group from the NT group. Yet, the present study was designed to find a feasibility of decoding individual pitches to set a basis for mentally producing any sequence of pitches through BCIs.

There is a slight possibility that the piano keyboard image and the horizontal movement of the indicator could affect the features when determining the ASR cutoff parameter. To address this, we examined decoding without the delta band reflecting the stimulus change at 2 Hz (ISI = 0.5 sec), and found no difference in decoding performance for all *K*-classes (Kruskal-Wallis test, p > 0.05). Moreover, the visual angle of the piano keyboard image was approximately 2.7°, which is difficult to be decoded even when using the artifact [44]. Thus, a possibility that decoding might be confounded by eye movements can be ruled out.

By verifying the feasibility of decoding seven pitches on the musical scale from human EEG, the realization of pitch imagery based BCI is surmised to be plausible. The feature extraction method proposed by this study may pave the way to discovering neural correlates of pitch imagery, although it needs more refinement through further investigations. However, it still requires an endeavor to enhance decoding performance to a level of guaranteeing practical BCI realization. To reach this, EEG feature reinforcement by training would be effective, as the neurofeedback training for MI-BCI improved corresponding EEG features [45]. The similar decoding performance of the MT and NT groups advocates that the future use can be applied to anyone regardless of their musical ability.

## V. Conclusion

This study revealed a feasibility to decode the imagined pitch on a musical scale from human EEG. We found spectral features that differentiated the multi-class pitches, represented the linearity of pitch height, and ruminated hemispheric differences. We achieved the performance of decoding the pitch imagery information from noninvasive brain signal, which could initiate the development of future pitch-imagery based BCIs for anyone who can represent pitch covertly heedless of the keen pitch sense.

