## Supplementary figures and images for "Decoding Imagined Musical Pitch from Human Scalp Electroencephalograms"

### FigS1

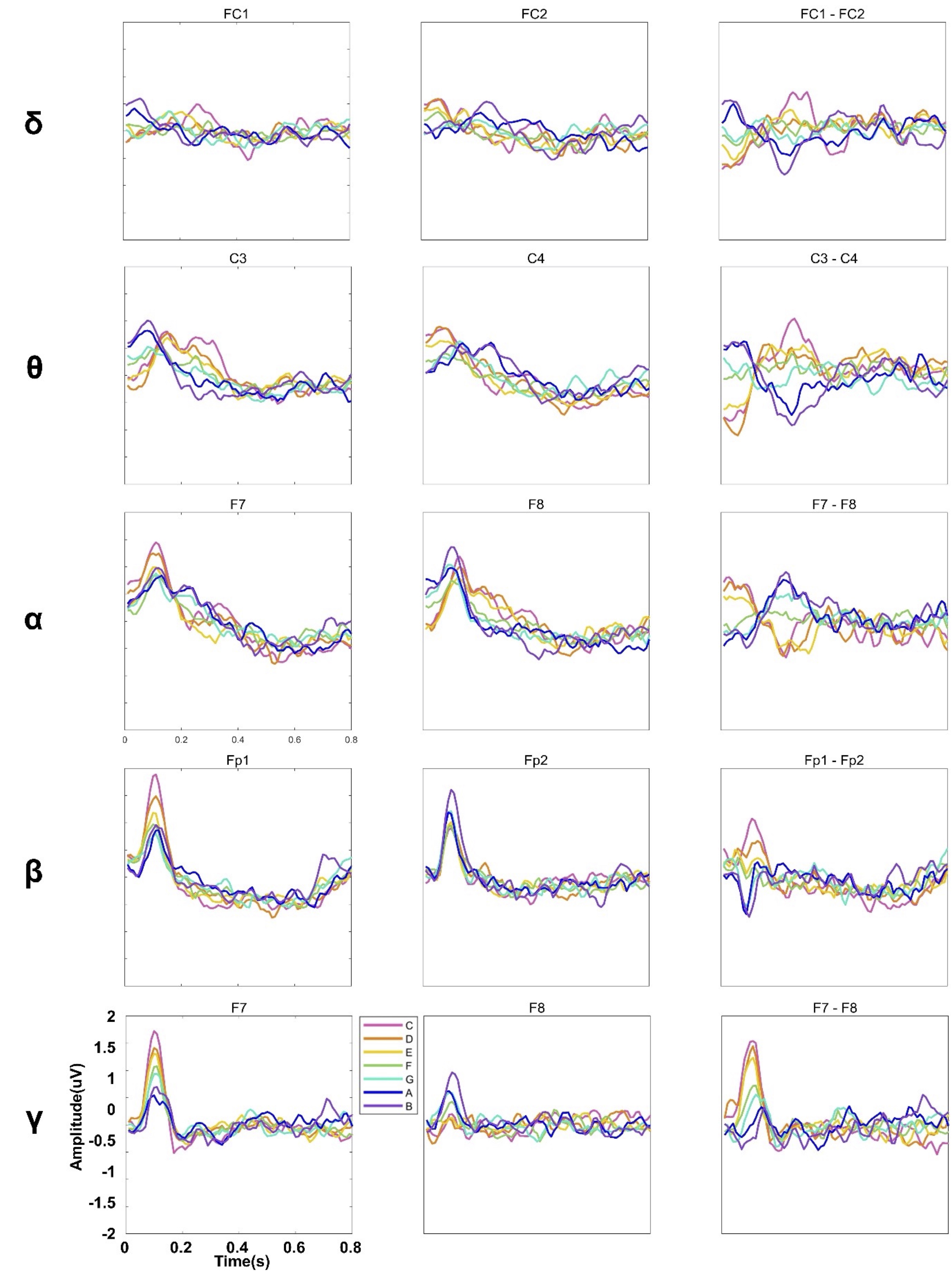
